# The loss of antioxidant activities impairs intestinal epithelium homeostasis by altering lipid metabolism

**DOI:** 10.1101/2023.03.09.531979

**Authors:** Javier Ramos-León, Concepción Valencia, Mariana Gutiérrez-Mariscal, David-David-Alejandro Rivera-Miranda, Celina García-Meléndrez, Luis Covarrubias

**Author notes:** Corresponding author: Luis Covarrubias, Av. Universidad 2001, Cuernavaca, Morelos, 62210, México, Web URL: http://www.ibt.unam.mx, /.

## Abstract

The increase in reactive oxygens species (ROS) with aging could be at the origin of many diseases of the elderly. Here we investigated the role of ROS in the renewal of the intestinal epithelium in mice lacking catalase (CAT) and/or nicotinamide nucleotide transhydrogenase (NNT) activities. *Cat^-/-^* mice have delayed intestinal epithelium renewal and were prone to develop necrotizing enterocolitis upon starvation. Interestingly, crypts lacking CAT showed fewer intestinal stem cells (ISC) and lower stem cell activity than wild-type, together with less LYS in Paneth cells. In contrast, crypts lacking NNT showed a similar number of ISCs and amount of LYS as wild-type but increased stem cell activity, which was also impaired by the loss of CAT. *Cat* deficiency caused fat accumulation in crypts, and a fall in the remarkable high amount of adipose triglyceride lipase (ATGL) in PCs. Supporting a role of ATGL in the regulation of ISC activity, its inhibition halt intestinal organoid development. These data suggest that the reduction of the intestine renewal capacity along aging originates from fatty acid metabolic alterations caused by peroxisomal ROS.

**Summary statement:** Mice with increased peroxisomal or mitochondrial reactive oxygen species develop intestinal phenotypes that are associated with aging and originate from a defective stem cell niche with impaired fatty acid metabolism.

## INTRODUCTION

Accumulation of cellular damage, emergence of senescent cells, inflammation and loss of regenerative capacity, all hallmarks of aging, are major challenges that tissues have to overcome to keep their functionality. In superior eukaryotes, cell replacement is a key process for tissue maintenance, particularly relevant when damages surpass the protection and repair mechanisms in cells. The intestinal epithelium is in constant replacement throughout life (every 3-5 days in mouse and 5-7 days in humans), being one of the tissues with the highest renewal rate, possibly due to the high cellular activity and damage during nutrient absorption and to the continuous exposure to the lumen adverse environment (Pentinmikko and Katajisto, 2020; Wang et al., 2021). Intestinal epithelium renewal is driven by a small population of adult stem cells (intestinal stem cells; ISCs) that reside in specialized and compartmentalized niches called crypts (Beumer and Clevers, 2021). The ISCs are the source of the different cell types that compose the intestinal epithelium, each with a specific role that give the functional characteristics to the intestine. The Paneth cells (PCs), also found in crypts, are interspersed between each ISCs and, in addition to their intestine protecting functions, have a determinant role in the stemness maintenance of ISCs by providing signals such as EGF, Notch ligands and Wnt (Gehart and Clevers, 2015; Sato et al., 2011) or, in extreme circumstances, by acquiring stem-like features (Schmitt et al., 2018). The loss of intestine regenerative capacity with aging has been linked to the exhaustion of ISCs, whereas the development of epithelial barrier leakiness, reduced antimicrobial protection and microbial dysbiosis, all alterations associated with the aged intestine, can directly relate to the loss of PC functions (Cray et al., 2021; Nalapareddy et al., 2022; Pentinmikko and Katajisto, 2020). In agreement with the relevant role of PCs in the generation and establishment of a fully functional intestine, the necrotizing enterocolitis syndrome, a common pathology of premature infants or newborns with low body weight, and the Crohn’s disease, a chronic inflammatory bowel pathology that affects young and old patients, are both associated with a loss in PC functions (Coutinho et al., 1998; Lueschow and McElroy, 2020).

Fatty acid (FA) oxidation plays a role in the maintenance and function of many stem cell populations including ISCs (Madsen et al., 2021). Accordingly, intestinal crypt cells have a large capacity to import FA (Chen et al., 2019), and metabolic interventions where FA metabolism is altered, such as fasting and high fat or ketogenic diets, have a significant impact on ISCs behavior (Cheng et al., 2019; Mana et al., 2021; Mihaylova et al., 2018; Wang et al., 2017). In addition, Cpt1-mediated FA oxidation in mitochondria of ISCs is a key requirement for their maintenance and induced intestinal epithelium renewal (Mihaylova et al., 2018), a mechanism that appears to be conserved in ISCs of Drosophila melanogaster (Singh et al., 2016). Downstream, HMGCS2, the limiting enzyme in the conversion of acetyl-CoA into ketone bodies, is also relevant for ISC maintenance. In this latter case, ketone bodies in ISC contribute to Notch signaling by inhibiting histone deacetylases (Cheng et al., 2019). Interestingly, the reduced number of ISCs in the intestine of aged mice correlates with a low FA oxidation activity (Mihaylova et al., 2018).

Very long chain fatty acids (VLCFA) are mainly catabolized in peroxisomes (Lodhi and Semenkovich, 2014). As a by-product of the peroxisomal β-oxidation of these VLCFAs, hydrogen peroxide (H_2_O_2_) is produced. Thus, several β-oxidation cycles from the starting VLCFA increase the levels of H_2_O_2_ and of several FA-derived metabolites in peroxisomes, including acyl-CoA derivatives that incorporate into mitochondria and contribute to establish the levels of acetyl-CoA and, under certain circumstances, also of ketone bodies. On the other hand, in mitochondria, through the trichloroacetic acid cycle, acetyl-CoA fuels the oxidative phosphorylation chain, a major source of reactive oxygen species (ROS; Martínez-Reyes and Chandel, 2020). Therefore, it is possible that, directly or indirectly, ROS produced in peroxisomes and mitochondria, in the course of lipid metabolism, contribute to aging in general and to the control of intestinal epithelium homeostasis in particular. The increase in peroxisomes in ISC after injury with the consequent promotion of intestine regeneration, observed in flies and mammals (Du et al., 2020), and the influence of oxidative phosphorylation and mitochondria-derived ROS in stem cell function in intestinal crypts (Rodríguez-Colman et al., 2017) agree with this possibility.

In tissues, the levels of ROS are determined by their production, frequently in association with metabolic activities such as those mentioned above, and also by the antioxidant capacity. Generally, the increase in ROS along aging is estimated from the accumulation of oxidative damage (Martin and Grotewiel, 2006; Luo et al., 2020), which in some instances has been correlated with an increase in antioxidant activities (Judge et al., 2005; Leutner et al., 2001; Ryan et al., 2008). However, the effects of ROS at an early aging stage, before oxidative damage accumulation, are not commonly appreciated as the origin of diseases of the elderly. Several antioxidant enzymes are compartmentalized in certain organelles where it is expected that they displayed a specific role. For instance, in peroxisomes, catalase (CAT) is the main antioxidant enzyme that prevents from H_2_O_2_ accumulation during FA β-oxidation (Lodhi and Semenkovich, 2014); thus, the impaired capacity of peroxisomes of aged cells to import catalase (Koepke et al., 2007; Terlecky et al., 2006), as well as other peroxisomal defects, could have significant pathological consequences (Cipolla and Lodhi, 2017; Uzor et al., 2020). In mitochondria, nicotinamide nucleotide transhydrogenase (NNT), an enzyme powered by the proton gradient generated by the respiratory chain, produces NADPH that, through its essential role in glutathione regeneration, provides an important antioxidant activity against the elevated production of ROS in active mitochondria (Rydström, 2006); mitochondrial failure has been propose at the bottom of many degenerative disease (Maurya et al., 2022) and, in particular, the lack of NNT causes glucocorticoid deficiency in mice and humans (Meimaridou et al., 2012).

Presently there is no evidence that the lack of CAT or NNT in mice causes an early in life general oxidative damage (Li et al., 2019; Pérez-Estrada et al., 2019), suggesting a mild and/or a restricted elevation in ROS in these conditions. Interestingly, however, a deficiency in either *Cat* or *Nnt* genes causes significant metabolic alterations of fatty acids (Fisher-Wellman et al., 2016; Pérez-Estrada et al., 2019; Rendina-Ruedy et al., 2015). In addition, the interaction between these two enzymes is revealed from the increased accumulation of fat in the liver and adipose tissue when both CAT and NNT are lacking (Heit et al., 2017; Shin et al., 2020). Here we report that the loss of CAT and/or NNT cause intestinal abnormalities, likely by a mechanism involving alterations in fatty acid metabolism, that parallel those found in the aged intestine and are causal of specific intestinal diseases.

## RESULTS

### *Cat* deficiency causes necrotizing enterocolitis and a delay in intestinal epithelium renewal

Fasting is a known condition that induces intestinal epithelium renewal. Phenotypically, as commonly observed due to fat accumulation, fasted Wt animals (*Nnt ^+/+^;Cat ^+/+^*) developed a pale liver (data not shown), and by 48 h of fasting, signs of intestinal necrosis (i.e., brownish aspect) were apparent (Fig. 1A, arrow). *Cat^-/-^* mice also developed a pale liver after fasting (data not shown) but, in contrast with Wt mice, a frequent incidence of necrotic intestine (i.e., black aspect) was noted from 24 h of fasting, and by 48 h, this phenotype became evident (Fig. 1A, arrows). Accordingly, only few apoptotic cells were detected in the intestine of Wt animals 24 h after fasting, whereas a significant emergence of apoptotic cells was evident in the intestine of *Cat^-/-^* mice in this same condition, mainly located within the crypt (Fig. 1B,C).

**Figure 1.**
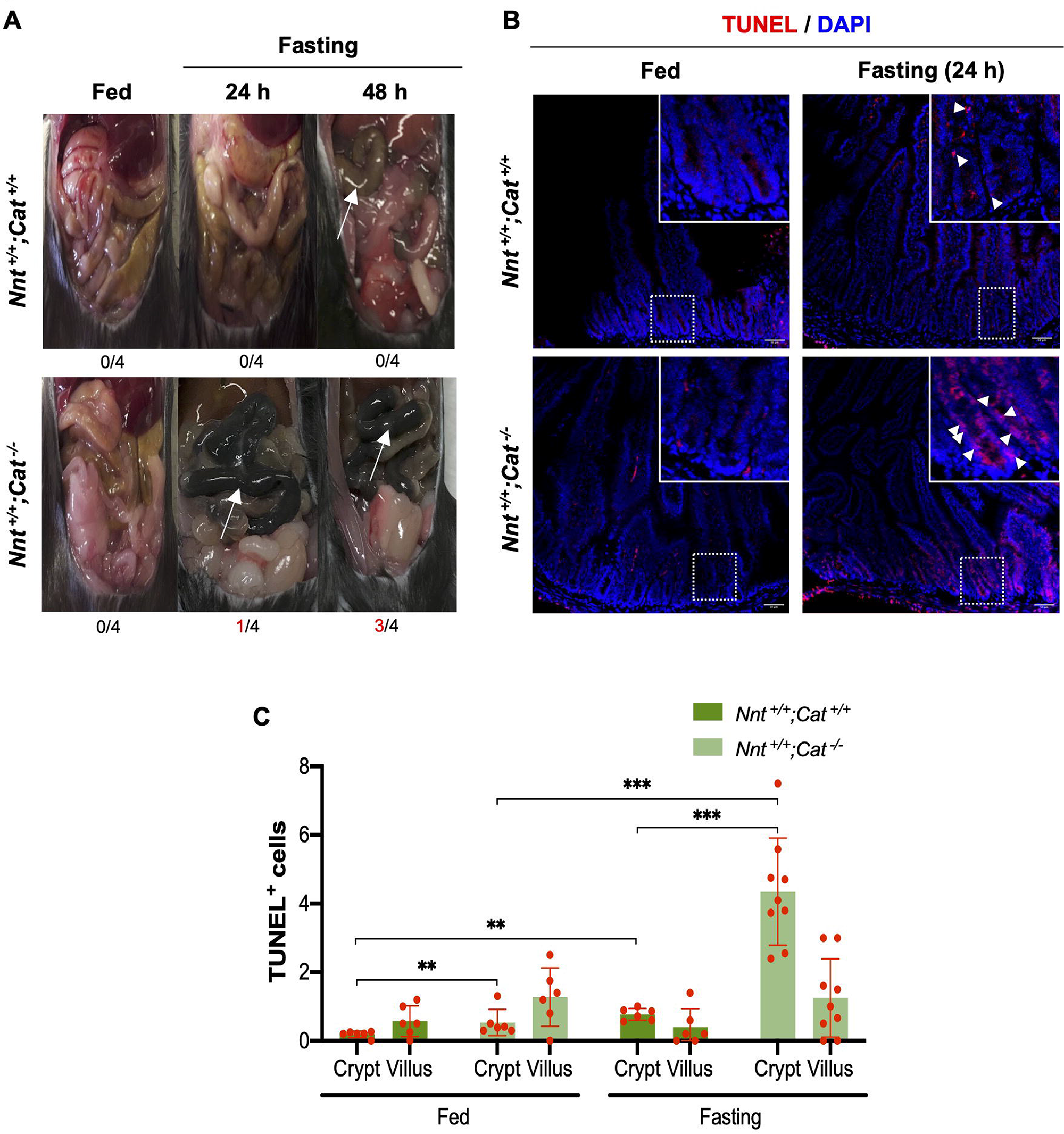
*Cat* is required for the maintenance of small intestine of mice under fasting conditions. (A) Gross appearance of small intestine of *Nnt ^+/+^;Cat ^+/+^*and *Nnt^+/+^;Cat^-/-^* mice in fed conditions and after fasting for 24 and 48 hours. Images of the most frequent phenotype in each condition for the genotypes indicated are shown; numbers below each image indicate the occurrence of mice with the necrotic intestine phenotype. Note the high incidence of necrotic intestine (arrows) in *Cat^-/-^* mice after fasting. (B,C) TUNEL assay staining and quantification in sections from duodenum of *Nnt ^+/+^;Cat ^+/+^*and *Nnt ^+/+^;Cat^-/-^* mice in fed and fasting (24 h) conditions. Representative images are shown (scale bar, 50 µm). Dying cells (arrowheads in inset) were mainly detected in crypts after fasting and more were found in the intestine of fasted *Cat^-/-^* mice. Data are shown as mean ± SD (*N* = 3), analyzed using a two-tailed unpaired Student’s *t*-test with significance *P* values of \*\**P* < 0.005 and \*\*\**P* < 0.0005.

The cell death observed could result from a limited capacity of the intestine of *Cat^-/-^* mice to carry out the characteristic epithelium renewal induced by fasting. In Wt mice, proliferating cells (Ki67+) were very abundant and mostly located at or near the crypt base compartment under fed conditions (Fig. S1A, *Nnt ^+/+^;Cat ^+/+^*) and, as previously demonstrated (Richmond et al., 2015; Wardill et al., 2020), fasting-induced renewal resulted in a significant decrease in these cells (Fig. S1A, Fasting). Interestingly, crypts of *Cat^-/-^* mice showed fewer proliferating cells than Wt under fed conditions (Fig. S1A, *Nnt ^+/+^;Cat^-/-^*), and no decrease due to fasting was observed (Fig. S1A, Fasting). Lack of NNT activity also caused a fall in proliferating cells in the crypt (Fig. S1A, *Nnt ^-/-^;Cat ^+/+^*) but, although the decrease in these cells due to fasting still occurred as in Wt, this effect was not observed when also CAT was lacking (Fig. S1A, *Nnt ^-/-^;Cat^-/-^*). Determination of *MKi67* mRNA levels in crypts was in agreement with the above observations showing downregulation in response to fasting in the presence but not in the absence of CAT (Fig. S1B). Thus, it is apparent that CAT is required for fasting-induced intestinal epithelium renewal.

To determine directly the role of CAT in the renewal of the intestinal epithelium, BrdU label-retaining cells were tracked after a single BrdU injection within a 4-day time window (Fig. 2A,B; (Parker et al., 2017)). BrdU incorporation in cells of the base was evident in Wt crypts 2 h after BrdU injection but, as Ki67^+^ cells, fewer were detected in the absence of either a peroxisomal (CAT) or a mitochondrial (NNT) antioxidant activities (Fig. 2C, 2 hpi). Conversely, particularly evident in crypts lacking NNT at this time, more BrdU-labeled cells in the transit amplification zone (TAZ) were detected than in Wt crypts (Fig. 2C, 2 hpi). As expected, 48 h after BrdU injection, it was rare to find Wt crypts with BrdU-labeled cells in the base region or TAZ and most BrdU label-retaining cells were found in the villi (Fig. 2C, 48 hpi, *Nnt ^+/+^;Cat ^+/+^*). Interestingly, BrdU- labeled cells in the base and TAZ were frequent in crypts lacking CAT, a characteristic that correlated with an increased number of BrdU label-retaining cells in the villi (Fig. 2C, 48 hpi, *Nnt ^+/+^;Cat^-/-^*). In contrast, a reduced number of BrdU label-retaining cells in the villi was observed in crypts lacking NNT in comparison with the number determined for Wt villi (Fig. 2C, 48 hpi, *Nnt ^+/+^;Cat ^+/+^* vs. *Nnt ^-/-^;Cat ^+/+^*). However, as in Wt crypts, the lack of CAT in the absence of NNT reverse this effect and an increased number of BrdU label-retaining cells were found in the villi (Fig. 2C, 48 hpi, *Nnt ^-/-^;Cat^-/-^*). Cell renewal in the intestinal tissue of mice of any genotype was completed by 96 h after BrdU injection since very few BrdU label-retaining cells were found at this time (Fig. 2C, 96 hpi). Therefore, the rate of removal of BrdU label-retaining cells was clearly slower in mice lacking CAT, independent of the presence or absence of NNT and, although less evident, lack of NNT alone showed a more rapid decay of BrdU labeled cells.

**Figure 2.**
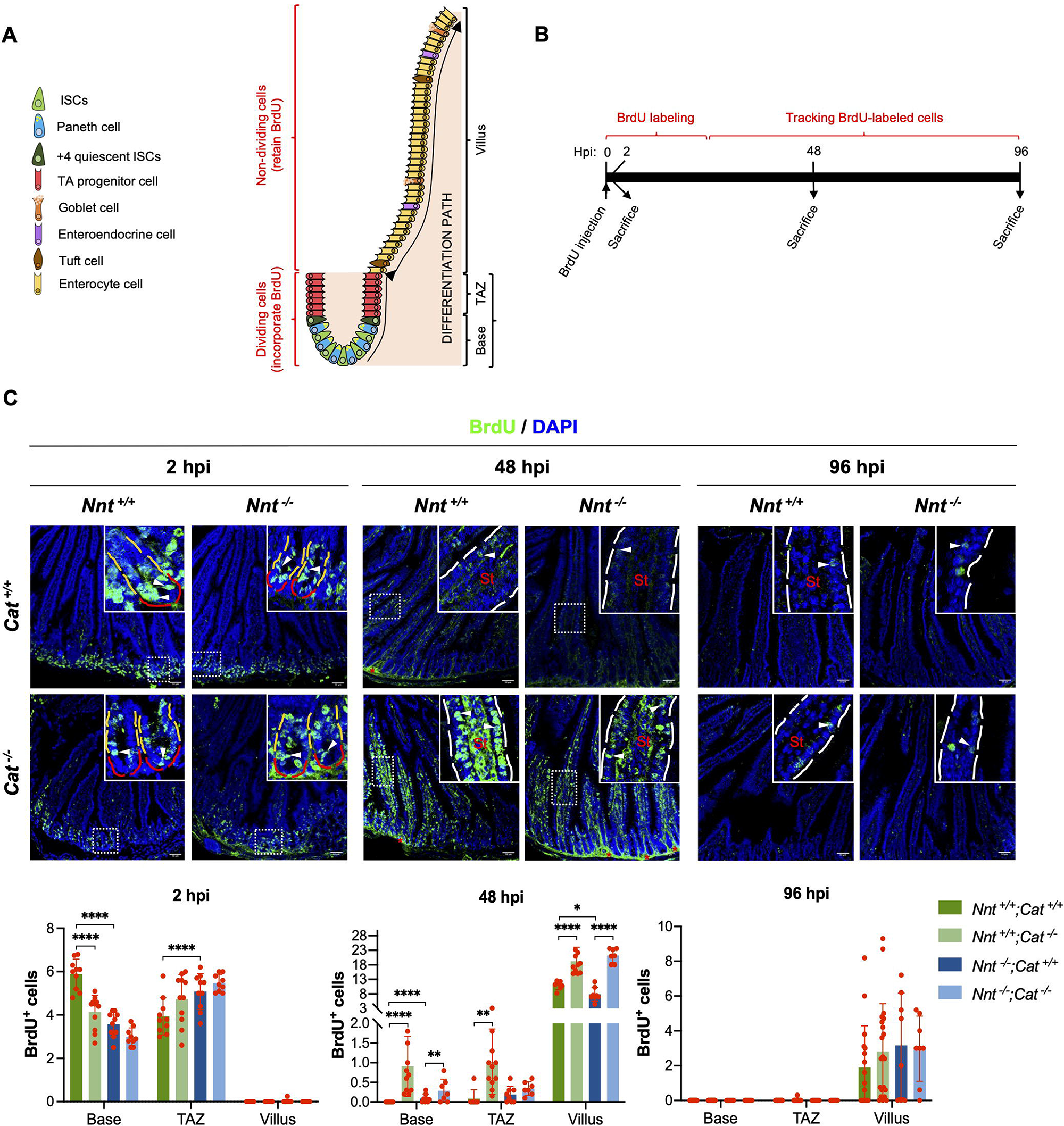
The loss of a functional *Cat* delays intestinal epithelium renewal. (A) The differentiation path of intestinal stem cells. Intestinal stem cells (ISC, green), in cooperation with Paneth cells (PC, blue), divide to maintain the stem cell niche which, upon activation, derive into transit amplifying cells (TAZ, red); these cells, with high proliferation activity, exit the cell cycle and give rise to the differentiated cells of the villus (enterocytes and Goblet, Tuft and enteroendocrine cells). (B) Protocol for BrdU incorporation and tracking of BrdU-labeled cells. After BrdU injection, mice were sacrificed at key stages of the intestinal epithelium renewal cycle. It is expected that within the first 2 hours after BrdU injection (2 hours post-injection, hpi) some cells of the crypt incorporate BrdU, and by 48 hpi, most cells retaining the BrdU label should be located in the villus, and completely removed after the epithelium renewal cycle ends (96 hpi). (C) Tracking of BrdU-labeled cells at 2, 48 and 96 hpi in sections from duodenum of *Nnt ^+/+^;Cat ^+/+^*, *Nnt ^+/+^;Cat ^-/-^, Nnt ^-/-^;Cat ^+/+^* and *Nnt ^-/-^;Cat^-/-^* mice under fed conditions. At 2 hpi, most BrdU-labeled cells were found in the base compartment (underlined in red in inset) or in the TAZ (underlined in yellow in inset); no BrdU-labeled cell was found in the villi at this time. At 48 hpi, most BrdU-labeled cells were found in the villi though, interestingly, a few were detected in crypts of *Nnt ^+/+^;Cat^-/-^* and *Nnt ^-/-^;Cat^-/-^* mice. Note the significant higher number of BrdU-labeled cells in the villi of mice lacking a functional *Cat* gene at 48 hpi. At 96 hpi, very few BrdU-labeled cells were found in the villi, an indication that the renewal cycle has ended. St, stroma; *, non- specific signal; scale bar, 50 µm. Dots in graph represent individual determinations in a single slice. Data are shown as mean ± SD (*N* = 5), analyzed using a two-tailed unpaired Student’s *t*-test with significance *P* values of \**P* < 0.05, \*\**P* < 0.005 and \*\*\*\**P* < 0.0001.

### Lack of CAT reduces ISCs in the intestinal crypt and goblet cells in the villi and affects PC functions

Under fed conditions, a histological analysis of intestine (duodenum) did not show major differences between Wt and *Cat^-/-^* mice (Fig. S2A), including similar number of crypts. Notably, as expected for a reduction in ISCs, a significant decline in the amount and mRNA levels of the ISC marker OLFM4 was detected in crypts of *Cat^-/-^* mice compared with crypts of Wt mice (Fig. 3A,C, *Nnt ^+/+^;Cat^-/-^* and vs. *Nnt ^+/+^;Cat ^+/+^*). This effect was not significant in crypts of *Nnt^-/-^* mice (Fig. 3A,C, *Nnt ^+/+^;Cat ^+/+^* vs. *Nnt ^-/-^;Cat ^+/+^*) but, again, the reduction in the amount and mRNA levels of OLFM4 due to the lack of CAT was observed in crypts of *Nnt ^-/-^;Cat^-/-^* double mutant mice (Fig. 3A,C, *Nnt ^-/-^;Cat ^+/+^* vs. *Nnt ^-/-^;Cat^-/-^*). The amount and mRNA levels of OLFM4 in crypts of Wt and *Nnt^-/-^* mice decreased after fasting (Fig. 3A,C, *Nnt ^+/+^;Cat ^+/+^* and *Nnt ^-/-^;Cat ^+/+^*, Fasting), as expected for the corresponding ISCs (Richmond et al., 2015) but, although this change was still apparent in the OLFM4 amount in crypts of fasted *Cat^-/-^* mice, particularly in those carrying the Wt *Nnt* allele (Fig. 3A, *Nnt ^+/+^;Cat^-/-^*), the *Olfm4* mRNA levels downregulation in response to fasting was not detected (Fig. 3C, Fasting), consistent with fewer ISCs in crypts of mice lacking CAT.

**Figure 3.**
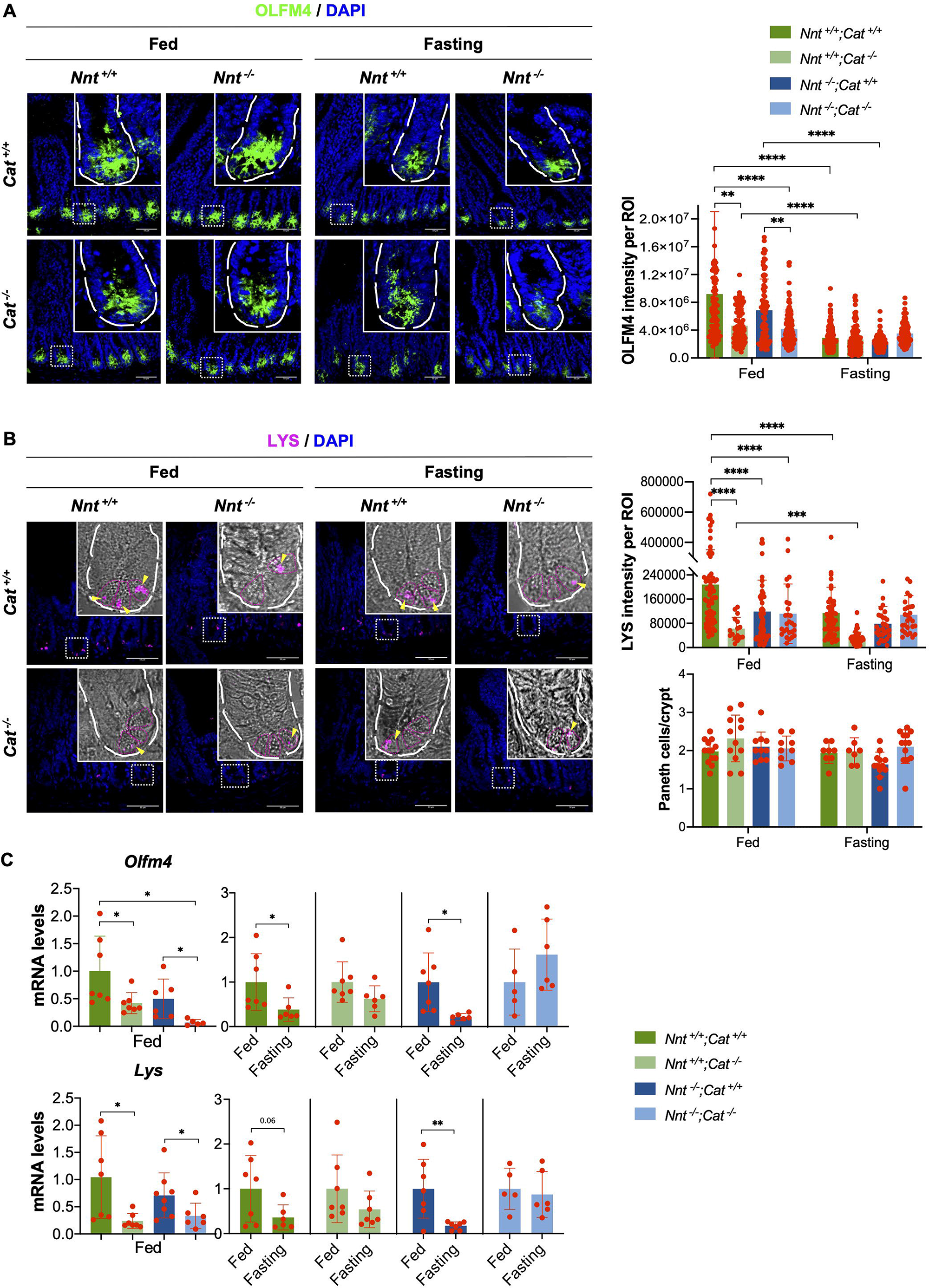
Lack of CAT reduces ISCs and LYS in the intestinal crypt and goblet cells in the villi. (A) OLFM4 immunostaining and quantification in sections from duodenum of *Nnt ^+/+^;Cat ^+/+^*, *Nnt ^+/+^;Cat ^-/-^, Nnt ^-/-^;Cat ^+/+^* and *Nnt ^-/-^;Cat^-/-^* mice in fed and fasting conditions. In representative images, a magnification of a single crypt is shown in insets. Observe that the amount of OLFM4 was reduced in the absence of CAT or upon fasting, independent of a functional *Nnt* gene. Dots in graph represent an individual crypt analyzed. Data are shown as mean ± SD (*N* = 5), analyzed using a two-tailed unpaired Student’s *t*-test with significance *P* values of \**P* < 0.05, \*\**P* < 0.005, \*\*\**P* < 0.0005 and \*\*\*\**P* < 0.0001. (B) LYS immunostaining and quantification in sections from duodenum of *Nnt ^+/+^;Cat ^+/+^*, *Nnt ^+/+^;Cat ^-/-^, Nnt ^-/-^;Cat ^+/+^* and *Nnt ^-/-^;Cat^-/-^* mice in fed and fasting conditions. Insets in representative images show a bright field and immunofluorescence merged images of a single crypt with delineated PCs (scale bar, 50 µm). Observe that, although the amount of LYS (arrowheads) was reduced in the absence of CAT and/or NNT, and further reduced upon fasting in the presence of a functional *Cat* gene, the number of PCs, as determined from their granulated morphology, did not change. Dots in graph represent determinations in a single crypt. Data are shown as mean ± SD (*N* = 5), analyzed using a two-tailed unpaired Student’s *t*-test with significance *P* values of \**P* < 0.05, \*\**P* < 0.005 and \*\*\**P* < 0.0005. (C) Relative *Olfm4* and *Lys* mRNA levels in crypts from *Nnt ^+/+^;Cat ^+/+^*, *Nnt ^+/+^;Cat ^-/-^, Nnt ^-/-^;Cat ^+/+^* and *Nnt ^-/-^;Cat^-/-^* mice in fed and fasting conditions. Observe that the mRNA levels of both genes decreased in the absence of CAT and a further reduction due to fasting only occurred in the presence of a functional *Cat* gene. Data are shown as mean ± SD (*N* = 5-7), analyzed using a two-tailed unpaired Student’s *t*-test with significance *P* values of \**P* < 0.05 and \*\**P* < 0.005.

A significant drop in the PC marker LYS was found in crypts of *Cat^-/-^* mice in comparison with equivalent samples of Wt mice (Fig. 3B, *Nnt ^+/+^;Cat ^+/+^* vs. *Nnt ^+/+^;Cat^-/-^*), observation that was in agreement with the reduction in *Lys* mRNA levels (Fig. 3C). Also, a moderate reduction in LYS and *Lys* mRNA levels was observed in crypts of *Nnt ^- /-^* mice, but this effect became evident in crypts of *Nnt ^-/-^;Cat^-/-^* double mutant mice (Fig. 3B,C, *Nnt ^+/+^;Cat ^+/+^* vs. *Nnt ^-/-^;Cat ^+/+^* and *Nnt ^-/-^;Cat^-/-^*). Fasting caused a decrease in LYS and *Lys* mRNA levels in crypts of Wt and *Nnt^-/-^* mice (Fig. 3B,C, *Nnt ^+/+^;Cat ^+/+^* and *Nnt ^-/-^;Cat ^+/+^*, Fasting), however, this phenomenon did not occur in crypts of mice lacking CAT (Fig. 3B,C, *Nnt ^+/+^;Cat^-/-^* and *Nnt ^-/-^;Cat^-/-^*, Fasting), possibly because the starting amount was already low. Therefore, CAT appears to substantially influence the production of LYS in PCs.

The reduction in LYS in the absence of CAT is a sign of loss of PC functions but could also relate to a decrease in the number of PCs. Determining the number of PCs based in cell morphology (i.e., granulated cells in crypt base) showed that fasting did not cause a decrease in granulated cells (i.e., putative PCs) in crypts of Wt mice, and similarly in crypts of mice lacking CAT or NNT (Fig. 3B). In addition, crypts of fed mice lacking CAT, despite showing a significant reduction in LYS, no decrease in granulated cells was observed in comparison with Wt, either in the presence or absence of NNT (Fig. 3B). These observations are in agreement with reports showing a decrease in LYS without altering PC number after fasting (Hodin et al., 2011) or in association with aging (Juricic et al., 2022). Therefore, the drop in LYS levels is revealing that the lack of CAT, rather than affecting the PC number, is hampering PC functions.

A reduction in ISCs and PCs could result in deregulated ISC differentiation. Actually, Notch1 and mitochondria, two mediators of crypt activity, define differentiation into Paneth and goblet cells (Ludikhuize et al., 2020). In agreement with altered ISC differentiation in mice lacking CAT, the intestinal villus of *Cat^-/-^* mice showed fewer goblet cells than the villus of Wt mice (Fig. S2B, *Nnt ^+/+^;Cat ^+/+^* vs. *Nnt ^+/+^;Cat^-/-^*). Also the lack of NNT alone caused a decrease in the number of goblet cells in villus compared with Wt, but a further decrease was not observed when CAT was also absent (Fig. S2B, *Nnt ^+/+^;Cat ^+/+^* vs. *Nnt ^-/-^;Cat ^+/+^*). No differences were found in the number of goblet cells near the crypts base compartment among mice with any *Cat* or *Nnt* genotype (Fig. S2B). These observations are in agreement with altered cell differentiation into goblet cells due to either the lack of CAT or NNT.

### *Cat* and *Nnt* deficiencies alter intestinal stem cell activity

The capacity of intestinal crypts to form organoids is an indirect determination of stem cell activity in crypts, starting from the formation of spherical organoids (SO) up to developing budding organoids (BO) containing crypt-like domains (Fig. 4A). Thus, as expected from the above results, crypts of *Cat^-/-^* mice formed fewer spherical (SO) and budding (BO) organoids in comparison with those of Wt mice (Fig. 4B,C, , *Nnt ^+/+^;Cat ^+/+^* vs. *Nnt ^+/+^;Cat^-/-^*); note that it is apparent that crypts of *Cat^-/-^* mice initiate organoid formation but degenerate at an early developmental stage (Fig 4B, arrows). In addition, among organoids with at least one domain, more domains were found in organoids derived from crypts of Wt mice than in those from crypts of mice lacking CAT (Fig. 4D, *Nnt ^+/+^;Cat ^+/+^* vs. *Nnt ^+/+^;Cat^-/-^*). Interestingly, a higher number of total organoids was obtained from crypts of *Nnt^-/-^* mice than from crypts of Wt mice (Fig. 4B,C, *Nnt ^+/+^;Cat ^+/+^* vs. *Nnt ^-/-^;Cat ^+/+^*), but the number of domains per organoid remained unchanged (Fig. 4 B,D), suggesting that proliferating cells in the TAZ and/or increased number of PCs are a sign of high stem cell activity that, as occurred after fasting (Mihaylova et al., 2018), correlated with an increased capacity of crypts to form organoids. This increased activity, however, was still abrogated by the lack of CAT resulting in fewer organoids, though it was not evident in the number of domains per organoid (Fig. 4B-D, *Nnt ^-/-^;Cat ^+/+^* vs. *Nnt ^-/-^;Cat^-/-^*). Therefore, CAT appears to be required to maintain the stem cell function in crypts, whereas NNT influences the activation of those stem cells.

**Figure 4.**
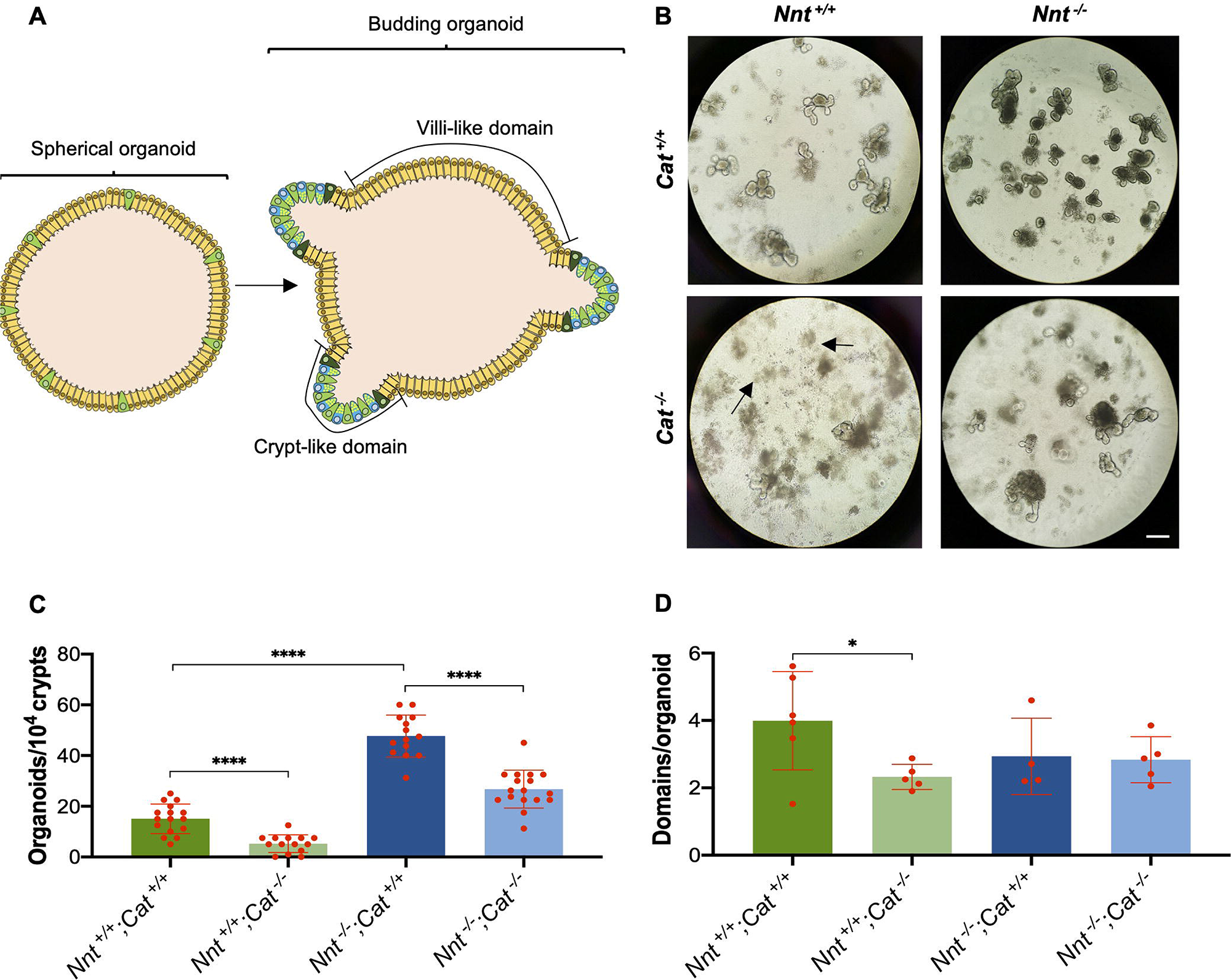
Crypt activity is altered by the lack of CAT and NNT. (A) Organoid formation to measure crypt activity. Crypts initially form spherical organoids that then develop into budding organoids; emergence of buds (i.e., crypt-like domains) is a sign of crypt activity. (B, C, D) Organoids generated from crypts of *Nnt ^+/+^;Cat ^+/+^*, *Nnt ^+/+^;Cat ^-/-^, Nnt ^-/-^;Cat ^+/+^* and *Nnt ^-/-^;Cat^-/-^* mice. Well-developed budding organoids were generated from crypts of *Nnt ^+/+^;Cat ^+/+^* and *Nnt ^-/-^;Cat ^+/+^* mice, but an increased number of them was obtained from crypts of the latter (B, C). The lack of CAT (*Nnt ^+/+^;Cat^-/-^*) caused organoid degeneration (arrows in B), and in the absence of NNT (*Nnt ^-/-^;Cat^-/-^*), the development of fewer budding organoids (B, C). A reduced number of crypt-like domains was only noted among the few organoids generated from crypts of *Nnt ^+/+^;Cat^-/-^* mice as compared with those originated from crypts of *Nnt ^+/+^;Cat ^+/+^* mice (D). Dots in C represent individual determinations in an organoid replicate culture, whereas in D is the average number of domains in organoids from an individual experiment. Data are shown as mean ± SD (*N* = 5), analyzed using a two-tailed unpaired Student’s *t*-test with significance *P* values of \**P* < 0.05 and \*\*\*\**P* < 0.0001.

### Lipid metabolism is the main target of antioxidant activities in the intestinal crypt

We previously showed that the lack of CAT alters lipid metabolism in the liver (Pérez-Estrada et al., 2019). In agreement with a similar effect in the intestine, accumulation of fat was observed in the intestine of mice lacking CAT, evident in the distal area of villi (Fig. S2C) but also noted in the crypt area (Fig. 5A). In intestinal crypts of mice lacking CAT, independent of the presence or absence of NNT, increased number of lipid droplets were found in many cells of the TAZ, and similar effect was also detected in several cells of the base, at least some in apparent ISC location (Fig. 5A). This observation could result from alterations in FA oxidation and ketogenesis, key processes required for ISC maintenance and/or differentiation (Cheng et al., 2019; Mihaylova et al., 2018). In particular, Mihaylova et al. (Mihaylova et al., 2018) showed that the expression of the gene encoding CPT1, the carnitine palmitoyltransferase essential for mitochondrial FA β-oxidation, increases in crypts in correlation with the intestinal tissue renewal induced by fasting. We did not detect a significant change in *Cpt1* expression levels in intestinal crypts of Wt mice upon fasting (Fig. 5B, Fasting, *Nn^+/+^;Cat ^+/+^*), even after 36 h of food removal (data not shown) but, interestingly, the expression levels did decrease in the intestinal crypts of mice lacking NNT in fed conditions (Fig. 5B, Fed, *Nnt ^+/+^;Cat ^+/+^* vs. *Nnt ^-/-^;Cat ^+/+^*). In contrast, expression levels of *Acox1*, encoding the rate limiting enzyme of straight chain FA β-oxidation in peroxisomes, were low in crypts of Wt mice and increased in response to fasting (Fig. 5B, Fed vs. Fasting, *Nnt ^+/+^;Cat ^+/+^*); this pattern was similar even in crypts without CAT (Fig. 5B, *Nnt ^+/+^;Cat^-/-^*). On the other hand, crypts lacking NNT showed increased *Acox1* expression levels under fed conditions in the presence but not in the absence of CAT, and in either case, an upregulation in response to fasting was not apparent (Fig. 5B, *Nnt ^-/-^;Cat ^+/+^* and *Nnt ^-/-^;Cat^-/-^*). As for *Acox1* under fed conditions, the expression levels of *Hmgcs2*, encoding an enzyme required for the synthesis of ketone bodies, were very low in crypts of *Nnt ^+/+^;Cat ^+/+^* and *Nnt ^+/+^;Cat^-/-^* mice and increased in those of *Nnt ^-/-^;Cat ^+/+^* but not in *Nnt ^-/-^;Cat^-/-^* mice (Fig. 5B). However, consistent with a previous report (Cheng et al., 2019), *Hmgcs2* expression levels (Fig. 5B) and amount (Fig. 5C) significantly increased in response to fasting, a phenomenon that was not affected by the lack of either CAT or NNT. Interestingly, HMGCS2 in crypts of mice lacking NNT under fed conditions was found in cells at the crypt base, that according with Cheng et al. (Cheng et al., 2019), should presumably be ISCs (Fig. 5C, *Nnt ^-/-^;Cat ^+/+^*, arrows); the number of these cells decreased when CAT was also absent (Fig. 5C, *Nnt ^-/-^;Cat^-/-^*, arrows) without any apparent change in the amount of enzyme due to the *Cat* or *Nnt* genotype or to feeding conditions (data not shown). These data suggest that the lack of NNT favors CAT-dependent peroxisomal lipolysis, and that peroxisomal FA β-oxidation and ketogenesis are NNT downstream activities that regulate ISC differentiation.

**Figure 5.**
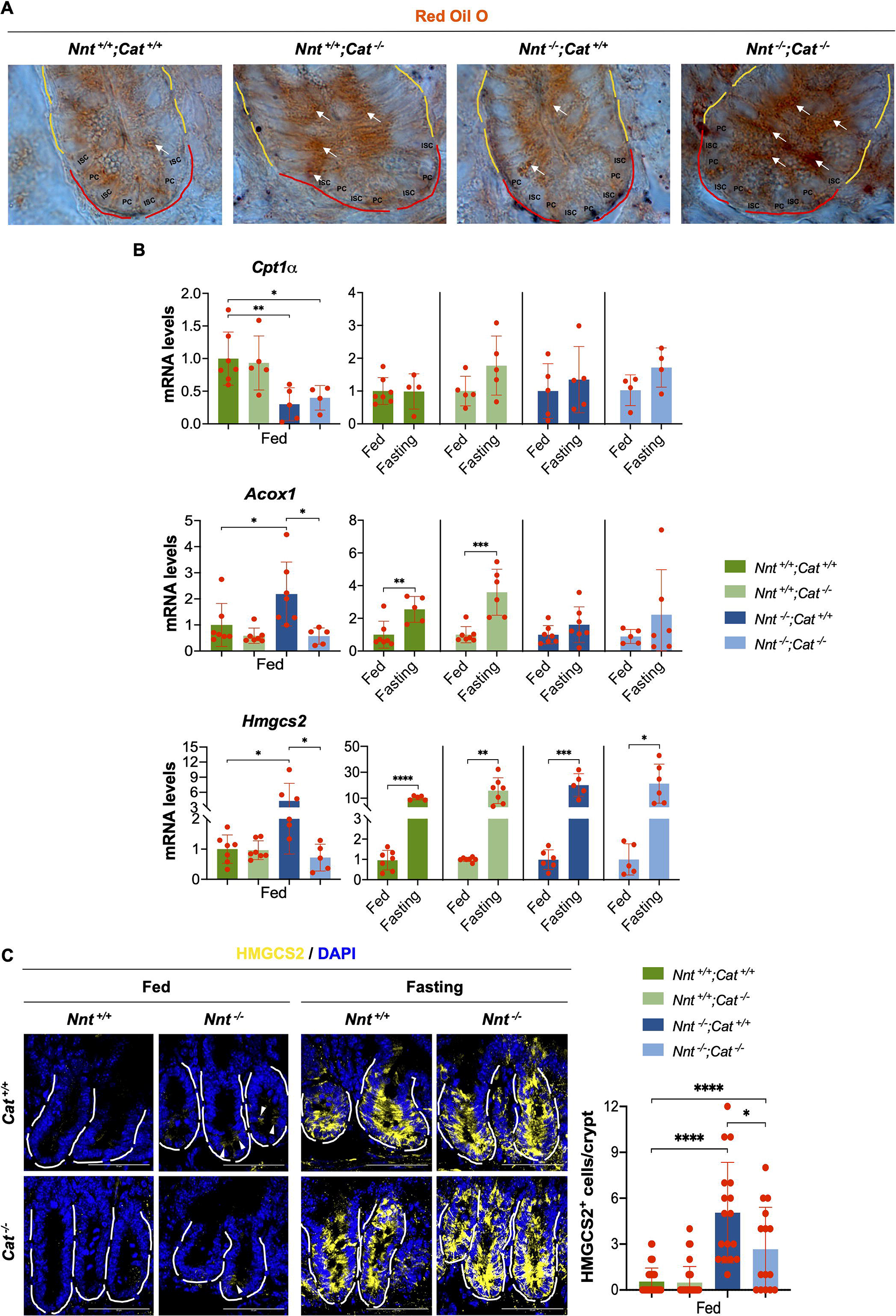
Lipolysis in the intestine appears altered by the loss of a functional *Cat* or *Nnt* gene. (A) Red Oil O staining of crypts in sections from duodenum of *Nnt ^+/+^;Cat ^+/+^*, *Nnt ^+/+^;Cat ^-/-^, Nnt ^-/-^;Cat ^+/+^*and *Nnt ^-/-^;Cat^-/-^* mice in fed conditions. Representative crypt (at least of 80% of total analyzed) for each genotype are shown (higher than . Lipid droplets (arrows) were detected in crypt cells, evident within the TAZ (underlined in yellow), but also some found in the base compartment (underlined in red), some in the apparent location of ISCs. (B) Relative mRNA levels of *Cpt1a, Acox1* and *Hmgcs2* in crypts from *Nnt ^+/+^;Cat ^+/+^*, *Nnt ^+/+^;Cat ^-/-^, Nnt ^-/-^;Cat ^+/+^*and *Nnt ^-/-^;Cat^-/-^* mice in fed and fasting conditions. *Cpt1a* mRNA levels were lower in crypts from mice lacking NNT (*Nnt^-/-^*) than in crypts from mice with the wild type allele (*Nnt ^+/+^*), whereas *Acox1* and *Hmgcs2* mRNA levels increased in the absence of NNT (*Nnt ^-/-^;Cat ^+/+^*) but decreased when CAT was lacking (*Nnt ^-/-^;Cat^-/-^*). In contrast with mRNA levels of *Cpt1a*, *Acox1* and *Hmgcs2* mRNA levels upregulated in response to fasting, though *Acox1* mRNA levels only increased in crypts of mice carrying the *Cat* wild type allele (*Cat ^+/+^*). Data are shown as mean ± SD (*N* = 5-7), analyzed using a two-tailed unpaired Student’s *t*-test with significance *P* values of \**P* < 0.05, \*\**P* < 0.005, \*\*\**P* < 0.0005 and \*\*\*\**P* < 0.0001. (C) HMGCS2 immunostaining and quantification in sections from duodenum of *Nnt ^+/+^;Cat ^+/+^*, *Nnt ^+/+^;Cat ^-/-^, Nnt ^-/-^;Cat ^+/+^*and *Nnt ^-/-^;Cat^-/-^* mice in fed and fasting conditions. Under fed conditions, HMGCS2^+^ cells (arrowheads) were only detected in crypts of mice lacking NNT (*Nnt^-/-^*) and fewer found when CAT was lacking (*Nnt ^-/-^;Cat^-/-^*). As expected, a large amount of HMGCS2 in crypts was detected in response to fasting, independent of the mouse *Cat* or *Nnt* genotype (scale bar, 50µm). Dots represent the number of positive cells or intensity in individual crypts analyzed. Data are shown as mean ± SD (*N* = 5), analyzed using a two-tailed unpaired Student’s *t*-test with significance *P* values of \**P* < 0.05, \*\**P* < 0.005, \*\*\**P* < 0.0005 and \*\*\*\**P* < 0.0001.

Peroxisome proliferation appears to be relevant for intestinal epithelium repair (Du et al., 2020). However, as determined by the amount of the peroxisomal protein PMP70 and expression levels of its gene, stimulation of intestinal epithelium renewal by fasting did not appear to cause any significant change in peroxisome number in crypts of Wt mice, though in the absence of CAT an increase was observed (Fig. S3A,B). Thus, no consistent changes in peroxisome amount due to the absence of CAT and/or NNT correlated with the renewal activity determined above. Coincident with a previous report in the liver (Oldford et al., 2019), the expression levels of *Cat* were increased in crypts lacking NNT, but the lack of any significant change in *Cat* mRNA levels in response to fasting did not correlate with crypt renewal activity (Fig. S3C). Interestingly, the *Acox1* expression levels described above did correlate with the crypt activity determined (Fig. 5B), suggesting that levels of peroxisomal β-oxidation activity does not associate with levels of CAT and can change without a corresponding similar increase in peroxisome number.

ACOX-mediated FA β-oxidation is an important source of H_2_O_2_ and CAT is a relevant enzyme that prevents its accumulation; therefore, lack of CAT could cause oxidative damage within the peroxisome and, thus, alter crypt activity. In order to determine the potential role of peroxisomal H_2_O_2_ in intestinal crypt activity, crypts were treated with an ACOX inhibitor (TDYA) one day after seeding for organoid formation (Fig. 6A). Using this protocol, ACOX inhibition did not have any significant effect on the capacity of crypts to form organoids (Fig. 6B,C), but an increase in the number of domains within formed organoids from crypts of Wt mice (Fig. 6B,D, *Nnt ^+/+^;Cat ^+/+^*) and a partial rescue of the reduced capacity of organoids from crypts of mice lacking CAT to develop domains were observed (Fig. 6B,D, *Nnt ^+/+^;Cat^-/-^*). Similar but mild effects of ACOX inhibition on organoid formation were detected when NNT was absent (Fig. 6B,D, *Nnt ^-/-^ ;Cat ^+/+^* and *Nnt ^-/-^;Cat^-/-^*). These observations support the notion that the effects shown in the absence of CAT are at least partially originated from an increase in peroxisomal H_2_O_2_ levels due to ACOX-mediated FA β-oxidation.

**Figure 6.**
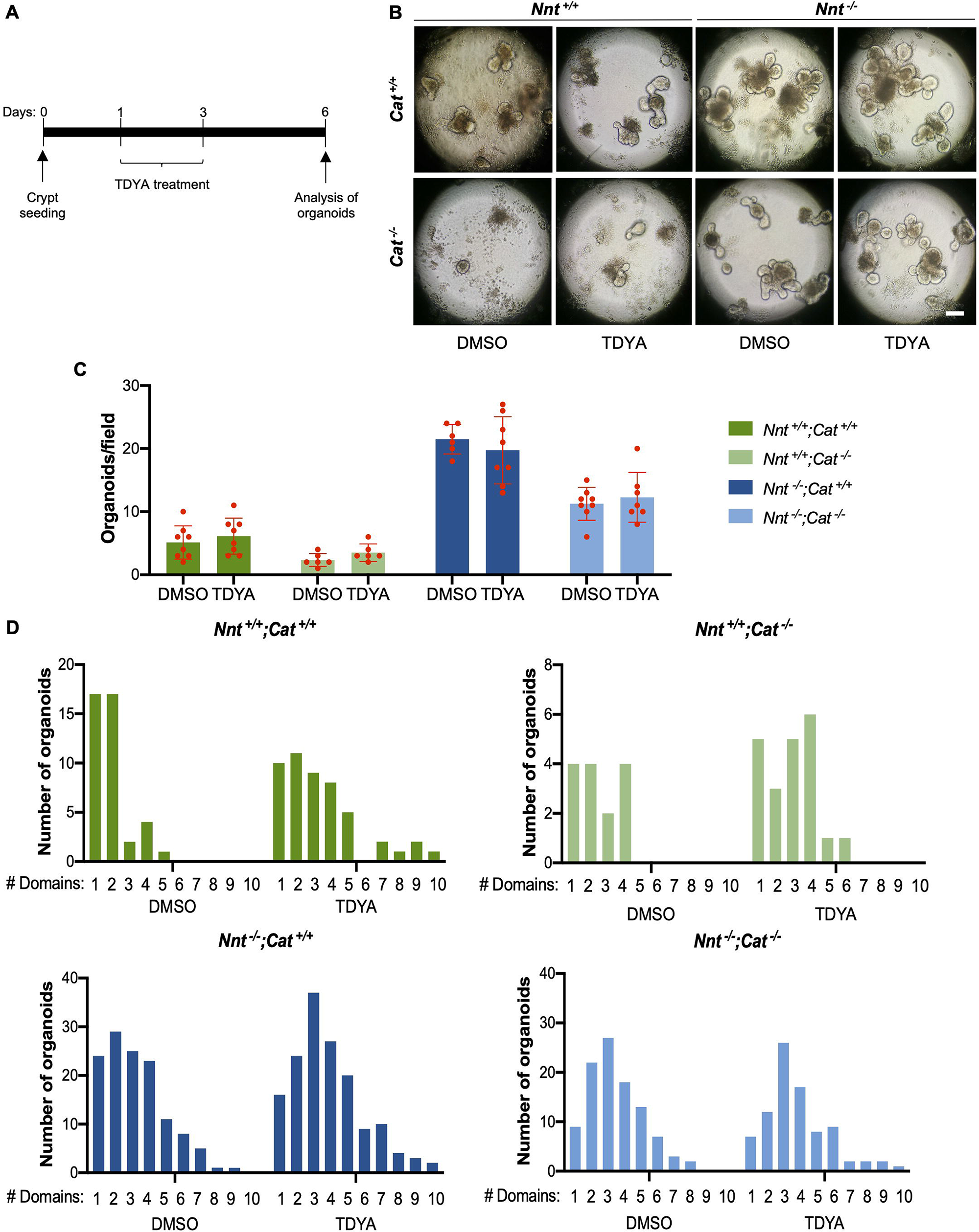
Inhibition of ACOX1 increases crypt activity. (A) Schematic representation of the protocol used for the treatment of intestinal organoids with the ACOX1 inhibitor TDYA. Development of organoids from isolated crypts was used to evaluate the influence of ACOX1 in crypt activity. TDYA ACOX1 inhibitor was added after spherical organoids were formed (i.e., one day after crypt seeding). TDYA treatment continued for two days, time at which the medium was replaced for medium without the inhibitor. Number of organoids as well as number of crypt-like domains per organoid were determined 6 days after crypt seeding. (B, C, D) Organoid formation and quantification of crypt-like domains per organoid generated from intestinal crypts of *Nnt ^+/+^;Cat ^+/+^*, *Nnt ^+/+^;Cat ^-/-^, Nnt ^-/-^;Cat ^+/+^*and *Nnt ^-/-^;Cat^-/-^* mice. Representative images of organoids generated are shown (B). The TDYA inhibitor did not cause any effect in the number of organoids obtained (B,C), but organoids with an increased number of crypt-like domains were observed in the presence of TDYA when compared with organoids generated in control conditions (DMSO) from crypts of any *Cat* or *Nnt* genotype (B,D). Determinations shown derive from 4 independent experiments. Dots in C represent individual determinations in an organoid replicate culture.

During fasting, peroxisomes can stimulate lipolysis by promoting the translocation of the adipose triglyceride lipase (ATGL) into lipid droplets (Kong et al., 2020). In addition, peroxisomal H_2_O_2_ can regulate lipolysis by directly controlling ATGL levels (Kong et al., 2020). Therefore, we explored the possibility that this mechanism is influencing crypt activity. ATGL was detected at low levels along the intestinal villi (Fig. S4A, asterisks) but, remarkably, cells with high levels of this enzyme were exclusively located within the crypt base (Fig. S4A, arrows), many of which were LYS^+^ cells (Fig. 7A). As expected from above observations regarding PCs, a reduced number of these ATGL^+^ cells were detected in crypts lacking CAT or NNT when compared with the number of these cells in Wt crypts (Fig. 7B). In addition to the reduction in the number of ATGL^+^ cells, the predicted reduction in the amount of ATGL per cell due to the lack of CAT was observed, though this phenomenon was not significant in the absence of NNT alone (Fig. 7B, *Nnt ^+/+^;Cat ^+/+^* vs. *Nnt ^+/+^;Cat^-/-^* and *Nnt ^-/-^;Cat ^+/+^*). Interestingly, although fasting did not cause a reduction in ATGL^+^ cells, ATGL amount did reduce in the presence of CAT (*Nnt ^+/+^;Cat ^+/+^* and *Nnt ^-/-^;Cat ^+/+^*) but not when CAT was lacking (*Nnt ^+/+^;Cat^-/-^* and *Nnt ^-/-^;Cat^-/-^*), in which case, an increase was apparent (Fig. 7B, Fasting). These latter observations could be due to the expected increase in H_2_O_2_ levels in association with fasting-induced lipolysis that should cause ATGL degradation, but the increase in crypts lacking CAT might originate, at least in part, from the induction of *de novo* ATGL synthesis, since upregulation of *Atgl* mRNA levels were detected in response to fasting, though only evident in the presence of NNT (Fig. 7C).

**Figure 7.**
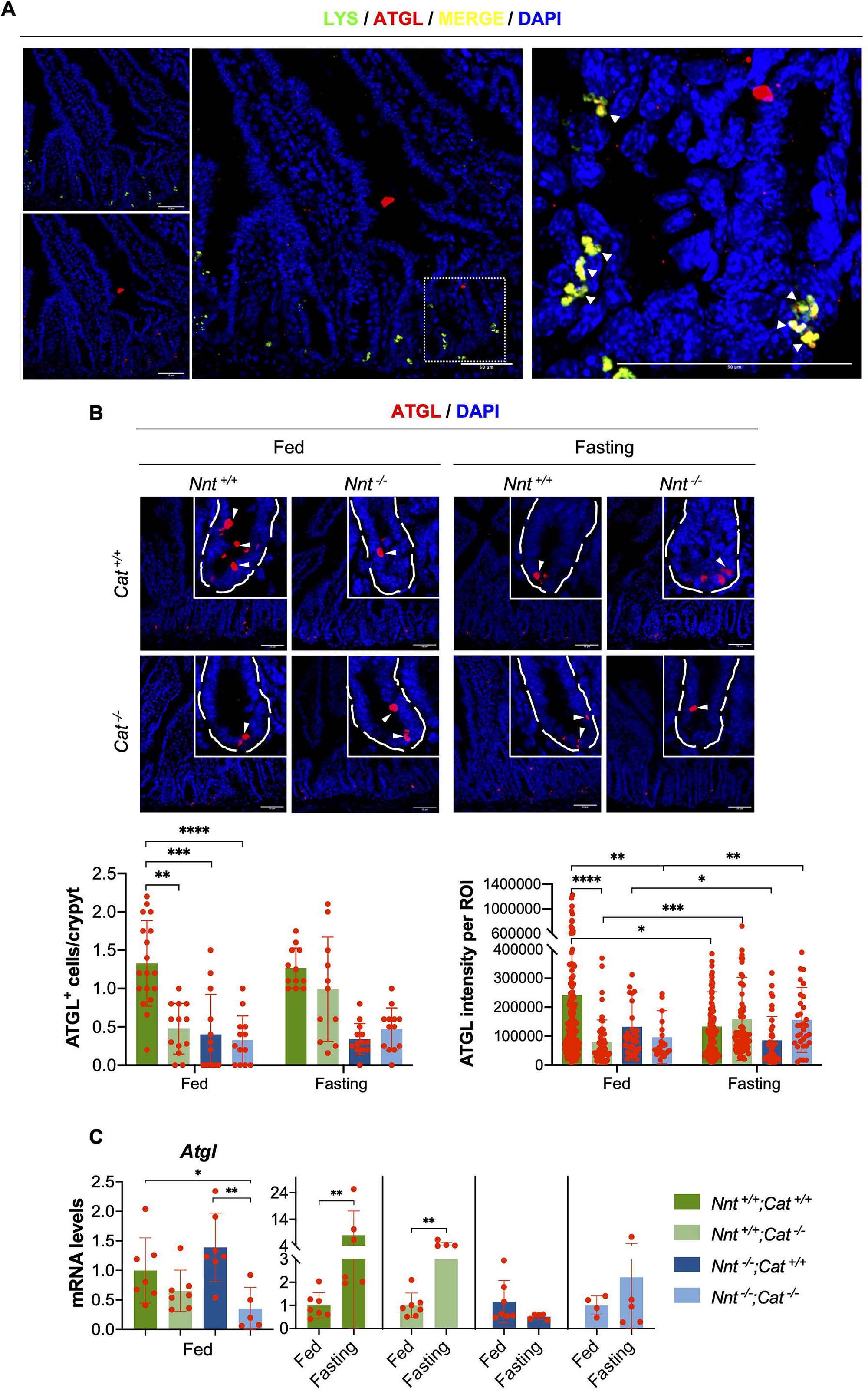
ATGL is localized in Paneth cells and reduced by the loss of a functional *Cat* or *Nnt* genes. (A) LYS and ATGL immunostaining in sections of duodenum of Wt mice. A double immunostaining was performed for LYS and ATGL proteins. Pictures are representative images of immunostained crypts. Cells with high ATGL levels were found in crypt base (see also B and Supplementary Fig. S3; scale bar, 50µm). Note that all LYS^+^ cells (green) colocalized with ATGL^+^ cells (red); arrows point LYS^+^/ATGL^+^ double stained cells. (B) ATGL immunostaining and quantification in sections from duodenum of *Nnt ^+/+^;Cat ^+/+^*, *Nnt ^+/+^;Cat ^-/-^, Nnt ^-/-^;Cat ^+/+^* and *Nnt ^-/-^;Cat^-/-^* mice in fed and fasting conditions. Representative images of crypts from mice of the genotypes indicated are shown. Observe that the lack of CAT (*Nnt ^+/+^;Cat^-/-^*) or NNT (*Nnt ^-/-^;Cat ^+/+^*) caused a decrease in the number of ATGL^+^ cells and ATGL amount per cell in comparison with crypts of Wt (*Nnt ^+/+^;Cat ^+/+^*) mice, but a response to fasting was only detected in the latter. Dots represent the number of positive cells or intensity in individual crypts analyzed. Data are shown as mean ± SD (*N* = 5), analyzed using a two-tailed unpaired Student’s *t*-test with significance *P* values of \**P* < 0.05, \*\**P* < 0.005, \*\*\**P* < 0.0005, \*\*\*\**P* < 0.0001. (C) Relative *Atgl* mRNA levels in crypts from *Nnt ^+/+^;Cat ^+/+^*, *Nnt ^+/+^;Cat ^-/-^, Nnt ^-/-^ ;Cat ^+/+^* and *Nnt ^-/-^;Cat^-/-^* mice in fed and fasting conditions. No major differences in *Atgl* mRNA levels among crypts with or without NNT or CAT were observed, but increased levels in response to fasting could be detected in crypts of mice carrying a wild type *Nnt* allele (*Nnt ^+/+^;Cat ^+/+^* and *Nnt ^+/+^;Cat^-/-^*). Data are shown as mean ± SD (*N* = 5-7), analyzed using a two-tailed unpaired Student’s *t*-test with significance *P* values of \**P* < 0.05, \*\**P* < 0.005.

A functional role of ATGL in crypt activity was tested in the organoid formation assay using an ATGL-specific inhibitor added one day after crypt seeding (CAY; Fig. 8A). The number of organoids derived from crypts of Wt mice was not affected by the ATGL inhibitor, but a significant reduction was noted in the absence of NNT (Fig. 8B,C, Total). In agreement with a relevant role of ATGL in the activity of intestinal crypts, ATGL inhibition at early stages (after day 1) of organoid formation resulted in a significant decrease in budding-organoid, an effect that was independent of whether organoids were derived from crypts of Wt or *Nnt^-/-^* mice (Fig. 8B,C, SO vs. BO). Therefore, these data suggest that peroxisomal H_2_O_2_ and mitochondrial ROS influences intestinal stem cell activity by regulating ATGL levels in PCs.

**Figure 8.**
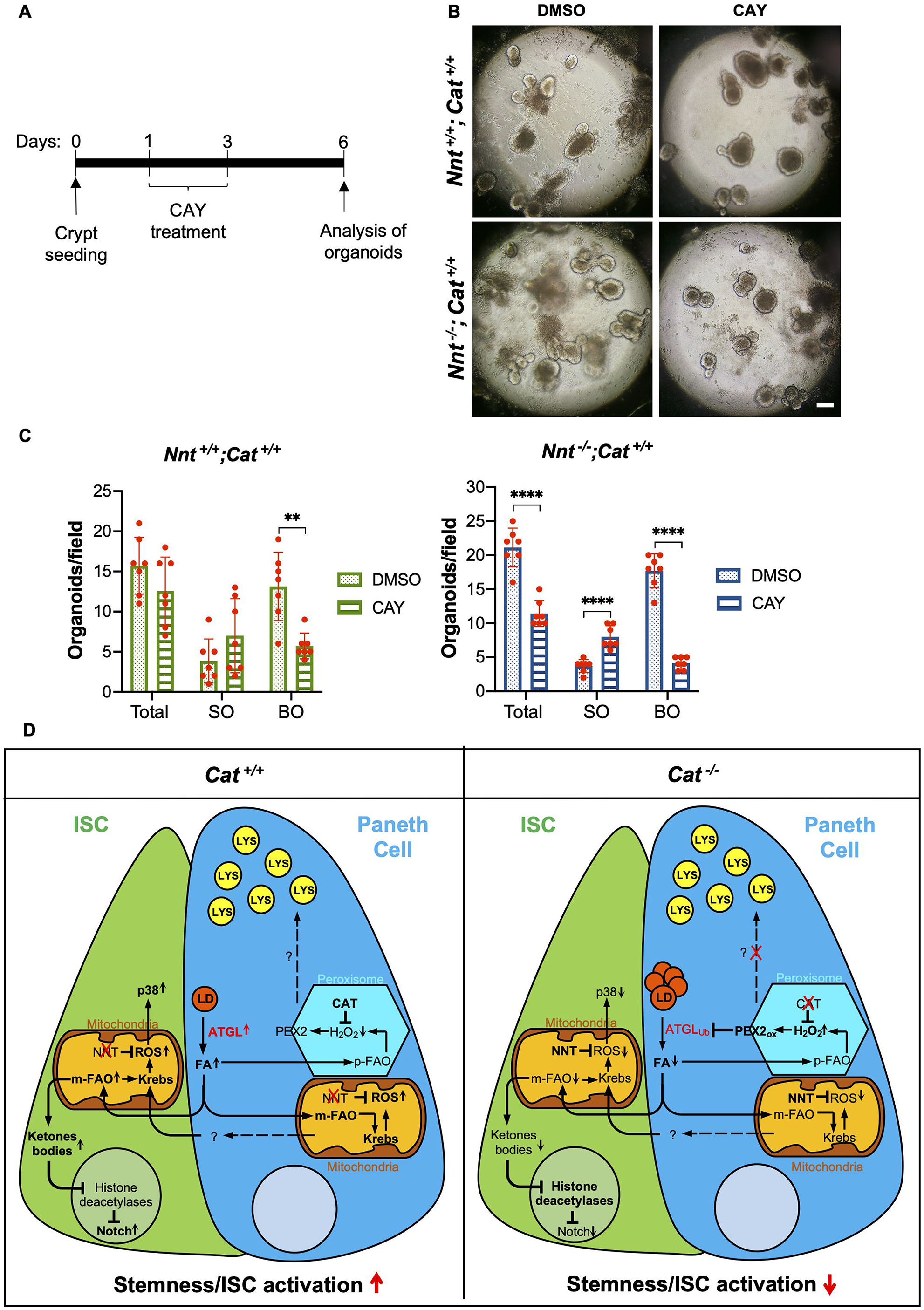
ATGL Inhibition reduces crypt activity. (A) Schematic representation of the protocol used for the treatment of intestinal organoids with the ATGL inhibitor CAY. Development of organoids from isolated crypts of *Nnt ^+/+^;Cat ^+/+^* and *Nnt ^-/-^;Cat ^+/+^* mice was used to evaluate the influence of ATGL in crypt activity. CAY ATGL inhibitor was added after spherical organoids were formed (i.e., one day after crypt seeding). CAY treatment continued for two days, time at which the medium was replaced for medium without the inhibitor. Number of spherical (SO) and budding (BO) organoids was determined 6 days after crypt seeding. (B, C) Organoid formation and quantification of SO and BO organoids generated from intestinal crypts of *Nnt ^+/+^;Cat ^+/+^* and *Nnt ^-/-^;Cat ^+/+^* mice. Representative images of organoids generated are shown (B). Lack of NNT caused a reduction in the total number of organoids as previously shown, but the CAY inhibitor prevented the formation of BO organoids, independent of the *Nnt* genotype. Dots in C represent individual determinations in an organoid replicate culture. Data are shown as mean ± SD (*N* = 5), analyzed using a two-tailed unpaired Student’s *t*-test with significance *P* values of \*\**P* < 0.005, \*\*\*\**P* < 0.0001.

## DISCUSSION

ROS are by-products of cellular metabolism. In particular, in mitochondria, high levels of ROS arise from the activity of the oxidative phosphorylation chain (OXPHOS; Murphy, 2009), and in the peroxisome, H_2_O_2_ is produced during the β-oxidation of, mainly, VLCFAs and branched FAs (Fransen and Lismont, 2019; Islinger et al., 2018). The large amount of ROS produced from this type of metabolic reactions have compelled cells to develop a variety of antioxidant activities that prevent cellular damage. However, since ROS also play a role in the regulation of cell signaling and of metabolic pathways (Lennicke and Cochemé, 2021), the level of antioxidant activities can also determine many stem cell activities, including proliferation and differentiation (Bigarella et al., 2014). Therefore, the progressive increase of ROS in the course of aging, in addition to favor cellular damage and thus tissue deterioration, can also alter fundamental cellular activities required to maintain a healthy tissue. In the present work, we show for the first time the significant influence of peroxisomal and mitochondrial ROS in the maintenance and activation of ISC and the overall homeostasis of the intestinal epithelium. Notably, the intestine of mice lacking CAT showed a reduction in: (i) ISCs and regenerative capacity, (ii) LYS amount in PCs, and (iii) goblet cells, all characteristics observed in the aged intestine but also associated with diseases affecting the intestine (e.g., Crohn disease, necrotizing enterocolitis; Demers-Mathieu, 2022; Du et al., 2020; Gebert et al., 2020; Juricic et al., 2022; Lee and Gulati, 2022; Pentinmikko and Katajisto, 2020; Wehkamp and Stange, 2020). In addition, we provide a strong support for a link between FA metabolism and the antioxidant activity, particularly that of CAT in peroxisomes, in the regulation of intestinal homeostasis. Our data suggest that ROS and FA metabolism are primary determinants of the loss of intestine regenerative capacity occurring in aged mammals and, although not studied here, also of other aging-associated dysfunctions that relate to a decline in PC and goblet cell functions.

### Mitochondrial antioxidant activity determines ISC differentiation

Different metabolic activities are required for the function of ISCs and PCs in the intestinal crypt. For instance, in contrast with PCs, mitochondria are very abundant in ISCs and mitochondrial activity is essential for their stem cell function (Rodríguez-Colman et al., 2017). Interestingly, the molecular components that determine this function are OXPHOS-derived ROS which, by stimulating p38 MAPK activity, induce crypt formation and differentiation of early intestinal epithelial cells into ISC and, also, appear to be required for maintaining ISC activity in mature crypts (Rodríguez-Colman et al., 2017). Therefore, it is expected that the lack of the antioxidant activity of NNT decreases the OXPHOS activity threshold at which ROS accumulate to signal ISC proliferation and/or differentiation. Here we show that, although no significant difference in the abundance of ISC in mice lacking NNT was detected, differentiation into proliferating TAZ was evident. This effect is in agreement with the proliferation promoting effect observed in the TAZ after crypt activation by fasting. Therefore, we propose that the lack of NNT changed the intestinal epithelium homeostasis dynamics such that crypt activation under basal condition, and possible also upon fasting, is more frequent than in Wt animals. This possibility is supported by the increased capacity of crypts lacking NNT to form organoids and the more rapid intestinal epithelium renewal observed in *Nnt* deficient mice.

Glycolysis-derived lactate in PCs seems to be the metabolite that feeds OXPHOS in ISC (Rodríguez-Colman et al., 2017). However, this could be the case under fed conditions, but under fasting, it has been shown that mitochondrial FA β-oxidation is activated in crypts, which can be revealed by the increased levels of *Cpt1* expression (Mihaylova et al., 2018). This response is coincident with the marked elevation in *Hmgcs2* expression upon fasting and the proposed role of ketone bodies in crypt activation (Cheng et al., 2019). Furthermore, genetically or pharmacologically blocking mitochondrial FA β-oxidation impairs crypt activity (Chen et al., 2019; Mihaylova et al., 2018), which contrast with the role of peroxisomal β-oxidation shown here (see below). We observed that fasting-induced *Cpt1* upregulation was a trend in crypts of mice lacking NNT, an effect that possibly relates to the lower *Cpt1* mRNA levels than Wt under fed conditions and, interestingly, is coincident with the higher levels of HMGCS2 and its mRNA. Therefore, it is apparent that the lack of NNT establishes a pre-activated condition in which FAs could be an additional source of metabolites that, cell autonomously or non-cell autonomously (see below), feed the OXPHOS in ISC. (Fig. 8D) The increased mRNA levels of *Acox1* and *Hmgcs2* in the absence of NNT suggest that these metabolites are produced in peroxisomes, and in mitochondria converted into ketone bodies, molecules capable to reinforce signaling pathways (e.g., Notch; (Cheng et al., 2019)) needed for efficient crypt activation. However, because it has been shown significant metabolic and ROS production alterations in liver mitochondria of C57BL/6J mice lacking active NNT (Oldford et al., 2019), the contribution of other metabolic pathways needs to be considered.

### Peroxisomes of Paneth cells regulate lipid metabolism and crypt activity

The peroxisome appears to be a key organelle in crypt activation. In conditions that stimulate intestinal repair such as inflammation, there is a significant increase in the number of peroxisomes (Du et al., 2020). However, this elevation in peroxisomes does not appear to accompany any induction of intestinal epithelium repair since, as shown here, peroxisomes levels in crypts of Wt and *Nnt^-/-^* mice after fasting did not show any significant change. Coincident with this observation, the levels of *Cat* mRNA also did not correlate with crypt activity. Therefore, peroxisome proliferation is not a general response in association with intestinal epithelium repair and, rather, depends on the status at which crypts are under basal conditions.

The contribution of peroxisomes to intestinal epithelium repair could be related to their activity rather than to their number. Peroxisomes carry out many biochemical reactions that produce ROS as by-products, being the ACOX-mediated FA β-oxidation the most important. Because CAT is mainly located in peroxisomes, its absence could result in peroxisomal rather than general protein damage. Accordingly, *Cat^-/-^* mice do not show significant oxidative damage in many tissues (Pérez-Estrada et al., 2019) but, as recently reported, the peroxisomal protein PEX2 is a target of peroxisomal H_2_O_2_, particularly evident in conditions that increase peroxisomal β-oxidation, causing its stabilization due to the formation of disulfide bonds (Ding et al., 2021). Interestingly, increased PEX2 levels mediates ATGL degradation by direct ubiquitylation in the proximity of lipid droplets (Ding et al., 2021). This mechanism explain why lipid metabolism is altered in the liver of *Cat^-/-^* mice, but with a differential outcome (i.e., increase or decrease in accumulation of fat) depending on the presence of NNT activity (Pérez-Estrada et al., 2019).

In the present work we found several evidence that suggest a key role of CAT as a regulator of lipid metabolism in the intestine. The negative role of peroxisomal FA β- oxidation in intestinal epithelium renewal is suggested from the increased capacity of crypt-derived intestinal organoids to develop domains when ACOX is inhibited, a condition that partially rescue the reduced capacity of organoids derived from crypts lacking CAT to form BO organoids. Remarkably, ATGL was found highly abundant in PCs, and as expected from an increase in H_2_O_2_ levels due to the lack of CAT, its amount was reduced in crypts of *Cat^-/-^* mice; the relevance of ATGL in crypt function is supported by the inability of intestinal organoids to develop domains when its activity is inhibited, a phenocopy of organoids from *Cat^-/-^* mice. Therefore, ATGL represents a key regulator of lipid metabolism in Paneth cells and a mediator of the negative effect of peroxisomal H_2_O_2_ on crypt activity (Fig. 8D). Notably, high levels of ATGL were found in colon cancer and cancer stem cells (Iftikhar et al., 2021), supporting the proliferation promoting effect of ATGL predicted from our observations.

Peroxisomal H_2_O_2_, by altering FA metabolism, could regulate crypt activity by different non-exclusive mechanisms: (1) A variety of lipolysis-derived FAs, some produced in the peroxisome, are major PPAR ligands (Lodhi and Semenkovich, 2014; Zechner et al., 2012). Particularly, H_2_O_2_ could impair PPARα ligand synthesis and, due to reduced PPARα activity and downstream FA oxidation, increase the amount of Notum, a Wnt inhibitor secreted by PCs that negatively controls crypt activity (Pentinmikko et al., 2019). Supporting this possibility, lack of ATGL downregulates PPARα target genes in the intestinal epithelium (Obrowsky et al., 2013). (2) FA synthesis influences ISC maintenance by sustaining PPARδ activity which, apparently, supports Wnt signaling in ISCs (Li et al., 2022). A role of H_2_O_2_ in this mechanism is in agreement with our previous observations suggesting that the lack of CAT causes a reduction in lipogenesis (Pérez-Estrada et al., 2019). (3) Ketone bodies in ISCs contribute to Notch signaling by inhibiting histone deacetylases (Cheng et al., 2019). Based in our present observations, we propose that, as lactate from PCs feeds mitochondrial activity in ISCs (Rodríguez-Colman et al., 2017), FAs released from PCs feeds mitochondrial FA oxidation and ketogenesis in ISCs (Fig. 8D), a mechanism that could depend on the organism metabolic condition and determined in PCs through the peroxisomal FA β-oxidation-ATGL interaction. With the present knowledge, it cannot be discarded that FAs processed in PCs are not terminally metabolized to Acetyl-CoA in their mitochondria. However, independent of this latter possibility, the non-cell autonomous function of ATGL remains as a mechanism that could significantly influence key processes in mitochondria of ISCs (i.e., OXPHOS and FA oxidation, Fig. 8D; (Mihaylova et al., 2018; Rodríguez-Colman et al., 2017)), and thus, intestinal epithelium renewal and repair.

Peroxisome-mitochondria interaction is a phenomenon that has attracted particular attention in recent years due to its relevance in metabolic regulation, especially lipolysis, and the significant implications expected in aging and disease development (Fransen et al., 2017; Pascual-Ahuir et al., 2017). Accordingly, preserving the mitochondrial network in *C. elegans* increases lifespan by a mechanism requiring FA oxidation and peroxisomal function (Weir et al., 2017). Relevant for the present study, increasing H_2_O_2_ (e.g., due to the lack of CAT) in peroxisomes of cultured cells causes an elevation in mitochondrial ROS levels but, interestingly, increasing mitochondrial ROS has no effect on peroxisomal redox (Ivashchenko et al., 2011). Thus, in crypts, lack of NNT might not directly affect peroxisomal functions, but lack of CAT could mediate its effect by perturbing both peroxisomal and mitochondrial functions. Thus, in addition to the role of H_2_O_2_ in the regulation of the non-cell autonomous mechanism mediated by ATGL referred above, H_2_O_2_ could play a cell autonomous function by influencing directly mitochondrial activity in ISCs and/or PCs (Fig. 8D). In either case, our present report establishes a framework for more detailed studies on cell autonomous and non-cell autonomous interactions between mitochondria and peroxisome in the intestinal crypt.

### Concluding remarks

The data presented suggest that peroxisomal CAT significantly influence lipid metabolism in PCs by regulating ATGL activity, whereas NNT mostly regulate fundamental mitochondrial functions in ISCs and, together, determine crypt activity. We show that deregulation of this mechanism results from diminished antioxidant activities that bring typical characteristics of the aged intestine, partly due to a reduced renewal capacity. In addition, although the depletion of LYS in PCs of crypts lacking CAT cannot be directly related to alterations in lipid metabolism, it is expected that this effect compromise PC fundamental functions required to maintain the integrity of the intestinal epithelium, a dysfunction that accompanies aging. As a corollary, significant implications are expected from our study in pathologies such as the necrotizing enterocolitis or inflammatory diseases such as the Crohn’s disease, in which PCs and/or the regenerative capacity are affected (Adolph et al., 2013; Deuring et al., 2014; Lueschow et al., 2018; Tremblay et al., 2016). Our data suggest that managing specific antioxidants and/or FA metabolism pathways could improve intestinal functions and translate into extended human health-span.

## MATERIALS AND METHODS

### Animal care and procedures

Wild type (Wt; *Nnt ^+/+^;Cat ^+/+^*) mice used here refer to those belonging to the C57BL/6N strain. The null *Cat* allele used is the one generated as described in Pérez-Estrada et al. (Pérez-Estrada et al., 2019), whereas the null *Nnt* allele is the one present in the C57BL/6J strain. These mice were backcrossed to generate the *Cat^-/-^* (in the “N” background), *Nnt^-/-^* (in the “J” background) and *Cat ^-/-^;Nnt^-/-^* (in the hybrid “NJ” background) mice used in the present study. The mouse genotype was determined by PCR performed on tail genomic DNA using specific oligonucleotide primers for the wild type and mutant *Cat* and *Nnt* alleles (Table S1); absence of CAT activity was occasionally verified by observing the lack of production of oxygen in the presence H_2_O_2_ from a small blood sample. For all protocols, 3-6 months mice of both sexes were used. They were maintained under controlled conditions of temperature and humidity, in a 12- hour light-dark cycle. Mice were fed with a standard diet (1218SX, Envigo, Indianapolis, IN, USA) with *ad libitum* access. For each treatment and condition, mice were previously synchronized to ensure that all were in a similar metabolic condition. For this, 24 hours prior to each protocol, mice were deprived of food for 12 hours and then fed for another 12 hours. After these 12 hours, mice in the fasted group were deprived of food while the fed group was sacrificed at this time. To evaluate cell turnover, mice were injected intraperitoneally with BrdU at a dose of 50 mg/kg and sacrificed 2, 48 and 96 hours after injection. Sacrifice was performed by cervical dislocation and the intestine was either collected and stored at -70°C or processed for crypt isolation. All animal procedures described here were approved by the Bioethical Committee of our Institute and rigorously meet international standards.

### Histochemistry and immunodetection procedures

The duodenum segment of small intestine was dissected and processed for cryostat sectioning (12 µm sections; CM1850 cryostat, Leica, Germany). Sections were stained with hematoxylin/eosin, Red Oil O or Alcian Blue according with standard protocols; samples were analyzed and photographed using a BX51 microscope (Olympus, Japan) coupled to a digital camera (Axiocam ERc 5s, Zeiss, Germany). Evaluation of proliferation in tissues using antibodies against Ki67 or BrdU (anti-human Ki67 and anti-BrdU monoclonal antibodies; Thermo-Fisher Scientific and BD BioSciences, USA, respectively) was performed as previously described {GarciaMelendrez:GKcY05su }. For determination of intestinal and metabolic markers, sections were washed for 30 min with PBS, then permeabilized with 0.3% Triton X-100 and incubated with an antigen retriever at 65°C for 30 min and at room temperature for 15 min, followed by a PBS wash and an incubation with a blocking solution (PBS 1X, 5% goat serum and 0.3% Triton X-100). After 60 min at room temperature, samples were incubated with the primary antibody overnight at 4°C and 1.5 hours at room temperature. Primary antibodies used were: anti-Lysozyme C coupled to Alexa Fluor 488 (1:1; Cat # sc- 518012, Santa Cruz Biotechnology, USA), anti-Olfm4 (1:100; Cat # 39141, Cell Signaling Technology, USA), anti-HGMCS2 (1:100; Cat # 20940, Cell Signaling Technology, USA), anti-ATGL (1:100; Cat # 2439, Cell Signaling Technology, USA) and anti-PMP70 (1:100; Cat # P0497, Sigma Aldrich, Germany). Samples were washed three times with PBS and, except for Lysozyme C, incubated with a secondary antibody for 2 hours at room temperature. Secondary antibodies used were: Goat anti-mouse IgG Alexa Fluor 488 (1:1000; Cat # A11001, Thermo-Fisher Scientific, USA), Goat anti-rabbit IgG Alexa Fluor 488 (1:200; Cat # A11008, Thermo-Fisher Scientific, USA), and Goat anti-rabbit IgG Alexa Fluor 594 (1:200; Cat # A11012, Thermo-Fisher Scientific, USA). After immunolabeling, all samples were counterstained with DAPI and mounted with ProLon Gold (Thermo-Fisher Scientific, USA) and photographed using an inverted confocal microscope (Olympus FV1000, Japan). For the analysis of crypt number, crypts were manually counted along the epithelium of each image every 10 µm distance in a minimum of 5 photographs (20X magnification) per mice. Paneth cells, Alcian-blue^+^, Ki67^+^, BrdU^+^ and ATGL^+^ cells were manually counted considering their morphology, location in crypt, transit amplification zone and villus. Cell death was determined by the TUNEL assay using the In Situ Cell Death Detection Kit Fluorescein (Cat # 11684795910; Roche, Germany); only a signal overlapping a DAPI-stained nucleus was considered positive. The quantification of OLFM4, LYS, PMP70, ATGL and HGMSCS2 signal was performed with the Image J software by first selecting a region of interest (ROI) and then measuring the relative intensity as described by Du *et al*. (Du et al., 2020).

### Isolation of small intestinal crypts

Mouse duodenum small intestine were washed with cold PBS, longitudinally opened, and once free of fat and capillary vessels, mucus was removed. The opened intestine then was cut into fragments and collected in a 15 ml tube with ice-cold PBS. The intestinal tissue fragments were washed by vigorous shaking for 30 seconds and fragments transferred to a new tube containing cold PBS, washed 3 additional times, and finally washed with PBS containing penicillin/streptomycin. The intestinal fragments treated as above were incubated in cold crypt dissociation solution (10 mM EDTA) with shaking for 10 min at 4°C. Finally, after incubation, the samples were shaken vigorously for 30 seconds, the epithelium fragments removed, and the crypt-enriched suspension centrifuged at 1,000 x g for 5 min. The pellet recovered was processed for RT-qPCR analysis or organoid culture after one wash with cold PBS or with cold DMEM-F12 w/HEPES, respectively.

### RT-qPCR for gene expression analysis in crypts

The PBS-washed suspension of crypt-enriched tissue, obtained as described above, was centrifuged at 1,000 x g for 5 min, and the pellet resuspended in cold PBS. This latter suspension was filtered through a 70-µm nylon mesh, and the crypts recovered by centrifugation at 1,000 x *g* for 5 min. The crypts were resuspended in RiboEx (GeneAll Biotecnology, Korea) for total RNA extraction following the manufactureŕs instructions. This RNA was converted into complementary DNA using the HyperScript^TM^ first strand synthesis kit (GeneAll, USA) with random hexameres as primers (Invitrogen, Thermo- Fisher Scientific, USA) or the Tribo^TM^ reverse transcription reaction kit (Tribioscience, USA). The quantitative PCR was performed in the Rotor-Gene Q (QIAGEN, Germany) using the KAPA SYBR^®^ FAST Universal qPCR Master Mix (2x; Kapa Biosystems, Inc., USA). Relative mRNA levels were normalized to *18S* expression levels via the ΔΔ method as previously reported (Pérez-Estrada et al., 2019). Primer sequences are listed in Table S1.

### Generation of intestinal organoids

The crypt-enriched suspension in cold DMEM-F12 w/HEPES (Gibco ThermoFisher, USA) was centrifuged at 1,000 x *g* for 5 min. The pellet was resuspended in cold DMEM-F12 w/HEPES containing 10 µM Y27632 (APExBIO, USA) and the resulting suspension was filtered through a 70-µm nylon mesh and centrifuged at 1,000 x *g* for 5 min. The obtained pellet was resuspended in 2 ml of cold DMEM-F12 w/HEPES and the number of crypts recovered determined. Four-thousand crypts were collected from this latter suspension, centrifuged, resuspended in 60 % Matrigel (ECM-GFR; Sigma-Aldrich, Merk, Germany) and seeded on a 37°C pre-warmed 48-wells plate (4,000 crypts/well); 20 min later, 250 µl of ENR organoid culture medium (DMEM/F12 w/o HEPES, Gibco; 1X Glutamax, Gibco; 10 mM HEPES, Sigma-Aldrich; 1X Penicillin/Streptomycin, Sigma-Aldrich; 1X MEM NEAA, Gibco; 1X Sodium Pyruvate, Gibco; 1X B27, Gibco; 1X N2, homemade; 1µM N-acetyl-L-cysteine, Sigma-Aldrich; 50 ng/ml EGF, Peprotech; 300 ng/ml Noggin, Peprotech; 200 ng/ml R-spondin-1, R&D) plus 10 µM Y-27632 (ENR-Y medium) was added. Twenty-four h later, the medium was replaced for ENR medium and changed every second day for 5-7 days. In some instances, the ACOX inhibitor 10,12-Tricosadiynoic acid (TDYA; Sigma-Aldrich, Merk, Germany) or the ATGL inhibitor [4-(5-methoxy-2-oxo-1,3,4-oxadiazol-3(2H)-yl)-2- methylphenyl]-carbamic acid, phenylmethyl ester (CAY10499; Cayman, USA), both dissolved in dimethyl sulfoxide (DMSO; Sigma-Aldrich, Merk, Germany) at the stock concentration of 500 µM and 2 mM, respectively, were added to the ENR medium (1 µM TDYA and 5 µM CAY final concentrations; for controls, equal volume as inhibitors of DMSO was added), and after 48 h of culture, the medium replaced for ENR medium alone for the following 5-7 days. Organoids were evaluated according with the number produced per crypts seeded and their morphology (i.e., spherical or budding organoids).

### Statistical analysis

Groups of mice with the same *Cat* or *Nnt* genotype and age were randomly assigned but always trying to keep a similar proportion of females and males; no evident sex-associated differences in the parameters measured were observed. The number of animals used in experiments for histological analyses were no less than 3 and generally between 5 and 6. Determinations in tissue slices were done at least 3 times in different regions of the same tissue fragment and from at least 5 mice (*N* are indicated in figures). For RNA analysis crypts derived from 5-7 mice and were individually processed but qPCR was performed in duplicate; in this case, to calculate the response to fasting, the average value of fed animals was set as “1”. For organoids, crypts derived from one mouse for each experiment (5 experiments), which was performed in duplicate. Data are shown as mean ± standard deviation (SD). All comparisons shown were between two groups applying a non-parametric two-tailed unpaired Student’s *t*- test; *P* < 0.05 is considered significant. Statistical analyses were performed with the GraphPad Prism software version 8.0b (La Jolla, CA, US).

## Acknowledgments

J.R.L. is a doctoral student from the Programa de Doctorado en Ciencias Biomédicas, Universidad Nacional Autónoma de México (UNAM) and has received CONACyT fellowship 620834. This study was supported by grants from the Programa de Apoyo a Proyectos de Investigación e Innovación Tecnológica (PAPIIT) of the Dirección General de Asuntos del Personal Académico (DGAPA) of the Universidad Nacional Autónoma de México (IN214219). We greatly acknowledge the support of the Animal Facility of the Instituto de Biotecnología along the whole study; particularly, the assistance by MVZ Graciela Cabeza Pérez, Oswaldo López Gutiérrez, Sergio González Trujillo and MVZ Elizabeth Mata Moreno is appreciated. We thank to the core members of the National Laboratory of Advanced Microscopy (LNMA), with a particular recognition to the technical support by Dr. Verónica Rojo.

## Author contributions

LC, led the research, designed experiments, analyzed the data, and wrote the manuscript. JR-L, designed and performed animal, histological and immunofluorescence experiments, analyzed the data, prepared the figures and contributed to the writing of the manuscript. CV, performed and analyzed organoid experiments. MG-M, performed and analyzed RT-qPCR experiments, and contributed to the statistical analyses. CG-M, contributed to the histological identification of Paneth cells and the immunodetection of lysozyme. D-AR-M, extracted total RNA from isolated crypts. All authors revised the final version of the manuscript.

## Notes

### Competing Interest Statement

The authors have declared no competing interest.

